# The lipid flippase ATP10B enables cellular lipid uptake under stress conditions

**DOI:** 10.1101/2023.06.01.543059

**Authors:** Rosanne Wouters, Igor Beletchi, Chris Van den Haute, Veerle Baekelandt, Shaun Martin, Jan Eggermont, Peter Vangheluwe

**Author notes:** equal contribution (Rosanne Wouters, Igor Beletchi). corresponding author Laboratory of Cellular Transport Systems ON1bis Herestraat 49 - box 802 3000 Leuven.

## Abstract

Pathogenic ATP10B variants have been described in patients with Parkinson’s disease and dementia with Lewy body disease, and we previously established ATP10B as a late endo-/lysosomal lipid flippase transporting both phosphatidylcholine (PC) and glucosylceramide (GluCer) from the lysosomal exoplasmic to cytoplasmic membrane leaflet. Since several other lipid flippases regulate cellular lipid uptake, we here examined whether also ATP10B impacts cellular lipid uptake. Transient co-expression of ATP10B with its obligatory subunit CDC50A stimulated the uptake of fluorescently (NBD-) labeled PC in HeLa cells. This uptake is dependent on the transport function of ATP10B, is impaired by disease-associated variants and appears specific for NBD-PC. Uptake of non-ATP10B substrates, such as NBD-sphingomyelin or NBD-phosphatidylethanolamine is not increased. Remarkably, in stable cell lines co-expressing ATP10B/CDC50A we only observed increased NBD-PC uptake following treatment with rotenone, a mitochondrial complex I inhibitor that induces transport-dependent ATP10B phenotypes. Conversely, Im95m and WM-115 cells with endogenous ATP10B expression, present a decreased NBD-PC uptake following ATP10B knockdown, an effect that is exacerbated under rotenone stress. Our data show that the endo-/lysosomal lipid flippase ATP10B contributes to cellular PC uptake under specific cell stress conditions.

## Introduction

Lipids are essential components of cellular membranes that play critical roles in maintaining cell and organelle structure and function. In addition, they determine the activity and localization of membrane proteins, operate as signaling molecules and function as an energy source (Van Meer, Voelker and Feigenson, 2008). The lipid composition in cells not only differs between membranes of various organelles, but also between the cytosolic and exoplasmic leaflet of a single membrane. This asymmetric distribution of lipids between the inner and outer leaflets of the membrane is an important feature of cellular membranes that needs to be carefully maintained to preserve the identity and function of a membrane (Kobayashi and Menon, 2018; Clarke, Hossain and Cao, 2020). However, the spontaneous movement of lipids across a membrane is generally a slow process, and an efficient directional movement of specific lipids between membrane leaflets depends on three classes of proteins: scramblases that move lipids bidirectionally, floppases moving lipids from the cytosolic to exoplasmic leaflet, and flippases, which move lipids from the exoplasmic to cytosolic direction (Pomorski and Menon, 2016).

Lipid flippases belong to the large family of P4-type transport ATPases, and several isoforms are involved in a broad-range of diseases, including neurodegeneration (Bull *et al*., 1998; Dhar *et al*., 2004; Levano *et al*., 2012; Onat *et al*., 2013; Yabas *et al*., 2014). P4-type ATPases typically contain 10 transmembrane domains for lipid translocation, and three cytosolic domains that couple lipid transport to ATP hydrolysis. During the catalytic cycle, lipid flippases spontaneously form an autophosphorylated intermediate on a critically conserved aspartic acid residue. Following the binding of a substrate lipid at the exoplasmic substrate binding site, transport often requires activation through a regulatory lipid or phosphorylation by a protein kinase, which relieves auto-inhibition by the N- or C-terminal extensions. This stimulates the substrate-dependent dephosphorylation reaction enabling lipid translocation (Andersen *et al*., 2016). Most P4-type ATPases require a regulatory subunit (CDC50A-C) for their proper functioning and localization (Takatsu *et al*., 2011; Naito *et al*., 2015) and mainly transport phospholipids, such as phosphatidylcholine (PC), phosphatidylserine (PS) and phosphatidylethanolamine (PE). More recently, members of the class V P4-type ATPase, isoforms ATP10A-D, have been described as lipid flippases that transport PC and/or the sphingolipid glucosylceramide (GluCer) (Naito *et al*., 2015; Roland *et al*., 2019; Martin *et al*., 2020).

We previously identified ATP10B as an endosomal lipid flippase that exports PC and GluCer out of late endo-/lysosomes that is mainly expressed in brain regions and the gastrointestinal tract (Martin *et al*., 2020). The specific subcellular localization of ATP10B relies on an endosomal targeting motif as well as on the interaction with the accessory subunit CDC50A that ensures ER-exit and proper subcellular localization (Takatsu *et al*., 2011; Naito *et al*., 2015). Compound heterozygous variants in ATP10B have been genetically associated with Parkinson’s disease (PD) and dementia with Lewy bodies (DLB), albeit with incomplete penetrance. This genetic link between ATP10B and PD/DLB has been the subject of discussion as it could not be recapitulated in some cohorts (Li *et al*., 2020; Smolders and Van Broeckhoven, 2020, 2021; Zhao *et al*., 2021). The identified pathogenic variants disturb ATP10B ATPase and lipid translocation activity, whereas ATP10B dysfunction affects lysosomal function and integrity, sensitizing neurons to mitochondrial oxidative stress and heavy metal exposure. ATP10B may operate in the lysosome together with the degradative enzyme glucocerebrosidase (GCase) to clear GluCer. GCase is encoded by GBA1, the most common genetic risk factor of PD (Do *et al*., 2019), which led to the hypothesis that ATP10B may represent a genetic risk factor of PD, although studies have so far been underpowered to confirm this.

The cellular lipid content is controlled by a coordination between lipid metabolism and uptake from the diet or bloodstream where lipids circulate as non-esterified free fatty acids (10%) or as esterified fatty acid components of triglycerides, phospholipids, and cholesteryl esters in lipoproteins (90%) (Abumrad *et al.,* 2021). The relative contribution of lipid uptake and metabolism to the lipid content in cells depends on several factors, such as the cell type, the type of lipid, the nutritional status, and the hormonal regulation. Adipocytes, hepatocytes and enterocytes display a high capacity for lipid uptake and storage, whereas muscle cells and neurons present a higher demand for lipid metabolism and utilization. Since the related plasma membrane (PM) isoforms ATP10A and ATP10D contribute to the cellular uptake of PC and/or GluCer (Naito *et al*., 2015; Roland *et al*., 2019), we here investigated whether ATP10B, as an intracellularly localized lipid flippase, may also promote the uptake of its substrate lipids into the cell. In overexpression and knockdown (KD) models, we found that ATP10B selectively stimulates cellular PC uptake.

## Materials and methods

### Cell culture and Transient Transfection

HeLa cells were routinely maintained in DMEM (Gibco, 41965-039) supplemented with 10% fetal bovine serum (FBS, Pan biotech 30-3306), non-essential amino acids (NEAA, Sigma, M7145-100ml), penicillin/streptomycin (Sigma, P4458-100ml), and if required relevant selection antibiotics (hygromycin 200 µg/ml ; InvivoGen, ant-hg-1 ; puromycin 2 µg/ml, InvivoGen, ant-pr-1). Im95m cells were routinely maintained in DMEM supplemented with 10% FBS; 10 µl/ml insulin (Merck, I9278), penicillin/streptomycin, and if required relevant selection antibiotics (blasticidine InvivoGen, ant-bl-1). WM115 cells were routinely maintained in DMEM supplemented with 10% FBS, NEAA, penicillin/streptomycin, sodium pyruvate (Gibco, 11360-039), and if required relevant selection antibiotics (blasticidin). All cell lines were cultured in an incubator with 5% CO_2_ at 37°C. When indicated cells were treated with rotenone (dissolved in dimethyl sulfoxide; Sigma-Aldrich, R8875). Generation of stable cell lines for ATP10B WT, ATP10B-GFP or ATP10B mutant expression and ATP10B KD with lentiviral vectors has been described in (Martin *et al*., 2020), and was performed by the Leuven Viral Vector Core using transfer plasmids for lentiviral vector production (Addgene, plasmid RRIDs: Addgene_171770, Addgene_171789, Addgene_204474, Addgene_204481, Addgene_171790, Addgene_171824, Addgene_204475, Addgene_171823, Addgene_204476; Addgene_204477, Addgene_204479, Addgene_204478, Addgene_204480) as described in dx.doi.org/10.17504/protocols.io.bw57pg9n.

For transient transfection, HeLa cells were co-transfected with pcDNA3.0 plasmids containing human CDC50A (NM_018247, N-terminal FLAG tag; RRID: Addgene_203694) and ATP10B variants (O94823.2, WT, RRID: Addgene_203695; catalytic mutations E210A and D433N, RRIDs: Addgene_203697 and Addgene_ 203696); and pathogenic variants R153X, V748L, E993A, G671R/N865K and I1038T, RRIDs: Addgene_203701, Addgene_203702, Addgene_203698, Addgene_203699, Addgene_203700) (Martin *et al*., 2020) using GeneJuice (Millipore, 70967-4) at a 3:1 ratio according to the manufacturer’s guidelines. After 24h experiments were performed.

### Microsomal isolation

Cells were harvested by detachment with TrypLE Express (Gibco, 12604-021), pelleted by centrifugation (5 min, 400 g_avg_, 4°C) and washed in phosphate buffered saline (PBS, Sigma, D8537-500ml), before being pelleted again. Cell pellets were lysed in RIPA lysis buffer (Thermo Scientific, 89900) supplemented with protease inhibitor (Sigmafast, Sigma, S8830-20TAB) for 30 min on ice. Lysates were centrifuged (10 min, 100 g_avg_, 4°C) to pellet and remove nuclear fraction. The resulting supernatant was centrifuged (10 min, 15 000 g_avg_, 4°C) to pellet the mitochondrial/lysosomal fraction. The resulting supernatant was centrifuged (35 min, 200 000 g_avg_, 4°C) to pellet the microsomal fraction, which was resuspended in sucrose (0.25 M). For subsequent Western blotting samples were prepared with 20 µg protein with 4x LDS loading buffer (NuPAGE, invitrogen, 2399420). A detailed protocol can be found on dx.doi.org/10.17504/protocols.io.rm7vzbw14vx1/v1.

### Western blot

Samples were loaded on 4-12% Bis/Tris gels (NuPage, Thermo Scientific, NP0321BOX). Gels were run in MOPS buffer (Life technologies, 1908733) for 35 min at 175 V. Proteins were transferred to 0.45 µm PVDF membrane (Immobilon-P, Thermo Scientific, 88518) for 60 min at 100 V. Membranes were blocked in blocking buffer (5% nonfat milk (Régilait, 0% fat) in TBS and 0.1% Tween20) for 1h at room temperature, before incubation with primary antibodies (ATP10B (Sigma-Aldrich Cat# HPA034574, RRID:AB_10669992), FLAG (Sigma-Aldrich Cat# F3165, RRID:AB_259529), GAPDH (Sigma-Aldrich Cat# G8795, RRID:AB_1078991), Na/K-ATPase (Abcam Cat# ab7671, RRID:AB_306023)) diluted in blocking buffer (4°C, overnight). The next day membranes were wash in TBS-T and incubated with peroxidase-conjugated secondary goat anti-rabbit or goat anti-mouse antibodies for 1 h at room temperature.

After final washing step in TBS-T immunodetection was performed on a Bio-Rad Chemidoc Imager with Pierce EC24L Immunoblotting substrate (Thermo Fisher Scientific, thermo scientific, 34580). Images were analyzed with Image Lab software (Biorad). A detailed protocol can be found on dx.doi.org/10.17504/protocols.io.ewov1oemklr2/v1.

### Lipid uptake

The lipid uptake assay was adapted from (Naito *et al*., 2015). Cells were harvested at 70% confluence by detachment with TrypLE Express). Cells were pelleted by centrifugation (5 min, 400 g_avg_, 4°C) and washed by PBS. After an additional centrifugation step (5 min, 400 g_avg_, 4°C), the cell pellet was resuspended in Hanks balances sodium solution (HBSS Gibco, 14170-112) and counted. Per condition 1*10^6^ cells were used in 500 µl final volume and equilibrated at 37°C for 15 min. An equal volume of 2 μM Nitrobenzoxadiazole (NBD)-labeled lipids (Avanti Polar lipids: NBD-PC (18:1-06:0 NBD PC, 810132C); NBD-lyso-PC (12:0 lyso NBD PC, 810128C); NBD-PS (18:1-06:0 NBD PS, 810194C); NBD-PE (18:1-06:0 NBD PE, 8105155C); NBD-SM (C6 NBD sphingomyelin, 810218C),NBD-GluCer (C6-NBD Glucosyl Ceramide, 810222C) were added to the cell suspension and incubated at 37°C for the required time point. At each subsequent time point, 200 μl of cell suspension was collected and mixed with 200 μl of ice-cold HBSS containing 5% fatty acid-free bovine serum albumin (BSA, Merck Millipore, 126575-10GM). To quench the fluorescence of non-internalized NBD-phospholipids, membrane-impermeable sodium dithionite (10 mM, Sigma-Aldrich) in 0.1 M Tris pH 10 (Sigma-Aldrich) was added 15 min prior to acquisition and mixed thoroughly. Next, cell samples were analyzed with an Attune NxT flow cytometer (Life Technologies) measuring the mean fluorescence intensity per cell of 10 000 events, after gating to exclude debris and doublets. A detailed protocol can be found on dx.doi.org/10.17504/protocols.io.14egn2pnpg5d/v1.

### Cell surface biotinylation

Cells were placed on ice and washed with ice-cold PBS. For 30 min cells were incubated on ice with PBS containing 2.5 mg/ml Sulfo-NHS-SS-biotin (Pierce). The biotinylation reaction was stopped by washing 3 times for 5 min with quenching solution (0.5% BSA and 100 mM glycine in PBS). Cells were lysed in RIPA buffer supplemented with protease inhibitor for 30 min on ice. Extracts were cleared by centrifugation (20 min, 14000 g_avg_, 4°C). Biotinylated proteins were isolated by incubation with immobilized Neutravidin beads (2 h rotating, 4°C). Input and bound proteins were processed for Western blotting with 4x LDS loading buffer. A detailed protocol can be found on dx.doi.org/10.17504/protocols.io.x54v9d9o4g3e/v1.

### Light microscopy and image analysis

Cells were seeded on coverslips and fixed with 4% paraformaldehyde for 20 min at room temperature and permeabilized with 0.1% Triton X-100 in PBS for 5 min. After blocking for 1 h with blocking buffer (PBS with 0.5% Tween20, 0.1% BSA, 0.2% FBS), cells were incubated with primary antibodies (anti-CD63, exbio, 11-343-C100, mouse; anti-GFP, abcam, ab13970, chicken) for 2 h at room temperature. Coverslips were washed with PBS-T and incubated 30 min with secondary antibodies (goat-anti-mouse-AlexaFluor647, goat-anti-chicken-AlexaFluor488). After washing with PBS-T, coverslips were incubated with DAPI, washed again and mounted using FluorSave. Images were acquired using an LSM780 confocal microscope (Zeiss) with a 10x or 40× objective. Colocalization analysis was performed with Fiji plugin Jacop. A detailed protocol can be found on dx.doi.org/10.17504/protocols.io.14egn2pypg5d/v1.

### Lipidomics

Lipidomics of HeLa samples, except for HexCer, were performed by Lipometrix according to following protocol: Lipid extraction: 700 μl of sample was mixed with 800 μl 1 N HCl:CH3OH 1:8 (v/v), 900 μl CHCl3 and 200 μg/ml of the antioxidant 2,6-di-tert-butyl-4-methylphenol (BHT; Sigma Aldrich). The organic fraction was evaporated using a Savant Speedvac spd111v (Thermo Fisher Scientific) at room temperature and the remaining lipid pellet was stored at - 20°C under argon. Mass spectrometry: Just before mass spectrometry analysis, lipid pellets were reconstituted in running solution (CH3OH:CHCl3:NH4OH; 90:10:1.25; v/v/v). Phospholipid species were analyzed by electrospray ionization tandem mass spectrometry (ESI-MS/MS) on a hybrid triple quadrupole/linear ion trap mass spectrometer (4000 QTRAP system; Applied Biosystems SCIEX) equipped with a TriVersa NanoMate (Advion Biosciences) robotic nanosource for automated sample injection and spraying. Phospholipid profiling was executed by (positive or negative) precursor ion or neutral loss scanning at a collision energy of 50 eV/45 eV, 35 eV, -35 eV, and -60 eV for precursor 184 (phosphatidylcholine (PC)), neutral loss 87 (phosphatidylserine (PS)) and precursor 241 (phosphatidylinositol (PI)), respectively. Phospholipid quantification was performed by multiple reactions monitoring (MRM), the transitions being based on the neutral losses or the typical product ions as described above. Lipid standards PC25:0, PC43:6, PI25:0, PI31:1, PI43:6, PS25:0, PS31:1, PS37:4 (Avanti Polar Lipids) were added based on the amount of [protein or DNA] in the original sample. Typically, a 3 min period of signal averaging was used for each spectrum. The data was corrected for isotope effects as described by (Liebisch *et al*., 2004).

Lipidomics of HexCer for HeLa samples was performed by the metabolomics facility at Washington University in St. Louis according to following protocol: Lipid extraction: Samples were homogenized in 1 ml of PBS. To homogenate sphingolipid internal standards were added, followed by protein precipitation with methanol, and collection of the supernatant. The supernatant was adjusted to 1 ml of 1:1 (v/v) methanol/water solution for HPLC/MS/MS. Instrumentation: Samples spiked with deuterated HexCer (d18:1 C18:0 d_35_) as internal standard were analyzed for glycosylceramides using the online trapping HPLC/MS/MS system. Calibration samples of the analytes, containing the same amount of internal standards, were also analyzed by the system. A Varian Metasil C18 column was used for analysis with a solvent gradient of 85% B (10 mM ammonium acetate (NH_4_OAc) and 1% formic acid) in 1:1 methanol/isopropanol and 15% A (10 mM NH_4_OAc and 1% formic acid in 3:7 acetonitrile/water) to 100% B in 5 min. Published positive Q_1_ and Q_3_ ions of these analytes were used for these analyses.

Lipidomics of WM115 samples were performed by Lipometrix according to following protocol: Lipid extraction: An amount of cells containing 10 μg of DNA was homogenized in 700 μL of water with a handheld sonicator and was mixed with 800 μl HCl(1M):CH3OH 1:8 (v/v), 900 μl CHCl3, 200 μg/ml of the antioxidant 2,6-di-tert-butyl-4-methylphenol (BHT; Sigma Aldrich), 3 μl of SPLASH® LIPIDOMIX® Mass Spec Standard (#330707, Avanti Polar Lipids), 3 μl of Ceramides and 3 μl of Hexosylceramides internal Standards (#5040167 and #5040398, AB SCIEX). After vortexing and centrifugation, the lower organic fraction was collected and evaporated using a Savant Speedvac spd111v (Thermo Fisher Scientific) at room temperature and the remaining lipid pellet was stored at - 20°C under argon. Mass spectrometry: Just before mass spectrometry analysis, lipid pellets were reconstituted in 100% ethanol. Lipid species were analyzed by liquid chromatography electrospray ionization tandem mass spectrometry (LC-ESI/MS/MS) on a Nexera X2 UHPLC system (Shimadzu) coupled with hybrid triple quadrupole/linear ion trap mass spectrometer (6500+ QTRAP system; AB SCIEX). Chromatographic separation was performed on a XBridge amide column (150 mm × 4.6 mm, 3.5 μm; Waters) maintained at 35°C using mobile phase A [1 mM ammonium acetate in water-acetonitrile 5:95 (v/v)] and mobile phase B [1 mM ammonium acetate in water-acetonitrile 50:50 (v/v)] in the following gradient: (0-6 min: 0% B → 6% B; 6-10 min: 6% B → 25% B; 10-11 min: 25% B → 98% B; 11-13 min: 98% B → 100% B; 13-19 min: 100% B; 19-24 min: 0% B) at a flow rate of 0.7 mL/min which was increased to 1.5 mL/min from 13 minutes onwards. PC, PI and PS were measured in negative ion mode by fatty acyl fragment ions. Lipid quantification was performed by scheduled multiple reactions monitoring (MRM), the transitions being based on the neutral losses or the typical product ions as described above. The instrument parameters were as follows: Curtain Gas = 35 psi; Collision Gas = 8 a.u. (medium); IonSpray Voltage = 5500 V and −4,500 V; Temperature = 550°C; Ion Source Gas 1 = 50 psi; Ion Source Gas 2 = 60 psi; Declustering Potential = 60 V and −80 V; Entrance Potential = 10 V and −10 V; Collision Cell Exit Potential = 15 V and −15 V. The following fatty acyl moieties were taken into account for the lipidomic analysis: 14:0, 14:1, 16:0, 16:1, 16:2, 18:0, 18:1, 18:2, 18:3, 20:0, 20:1, 20:2, 20:3, 20:4, 20:5, 22:0, 22:1, 22:2, 22:4, 22:5 and 22:6. Data Analysis: Peak integration was performed with the MultiQuant^TM^ software version 3.0.3. Lipid species signals were corrected for isotopic contributions (calculated with Python Molmass 2023.8.30; DOI: 10.5281/zenodo.7135495) and were quantified based on internal standard signals and adheres to the guidelines of the Lipidomics Standards Initiative (LSI) (level 2 type quantification as defined by the LSI).

### Data Access and Statistics

All data is available and can be accessed at Zenodo (DOI: 10.5281/zenodo.10044967). Statistics were performed and graphs were generated with Graphpad Prism software (version 9.4.1). Data are shown as mean ± SEM of a minimum of 3 independent experiments. Statistical test for multiple comparisons was an ordinary one-way ANOVA with Dunnett’s post-hoc pair-wise multiple comparisons test, for single comparisons was unpaired t test. Significance was set at *P<0.05, **P<0.01, ***P<0.001, and ****P<0.0001.

## Results

### Overexpression of ATP10B stimulates the cellular uptake of NBD-PC

To monitor the impact of ATP10B on cellular PC uptake we first turned to HeLa cells transiently transfected to co-express ATP10B and CDC50A. Several controls were included to study the lipid uptake caused directly by the lipid flippase activity of ATP10B: cells transfected with (i) only CDC50A (referred to as CDC50A), (ii) a catalytically dead mutant of ATP10B to control for non-catalytic functions of ATP10B, such as E210A or D433N, in which either Glu210 in the TGE dephosphorylation motif or Asp433 in the DKT autophosphorylation motif have been mutated, respectively, and (iii) five different pathogenic protein variants (R143X, G671T/N865K, V748L, E993A, I1038T) distributed across the catalytic domains (Fig. 1A) that abolish ATPase activity (Martin *et al*., 2020). Western blotting (Fig. 1B) after co-transfection confirmed that all cell lines expressed comparable levels of CDC50A and ATP10B, except the ATP10B truncation variant R153X, as previously reported (Martin *et al*., 2020).

**Figure 1:**
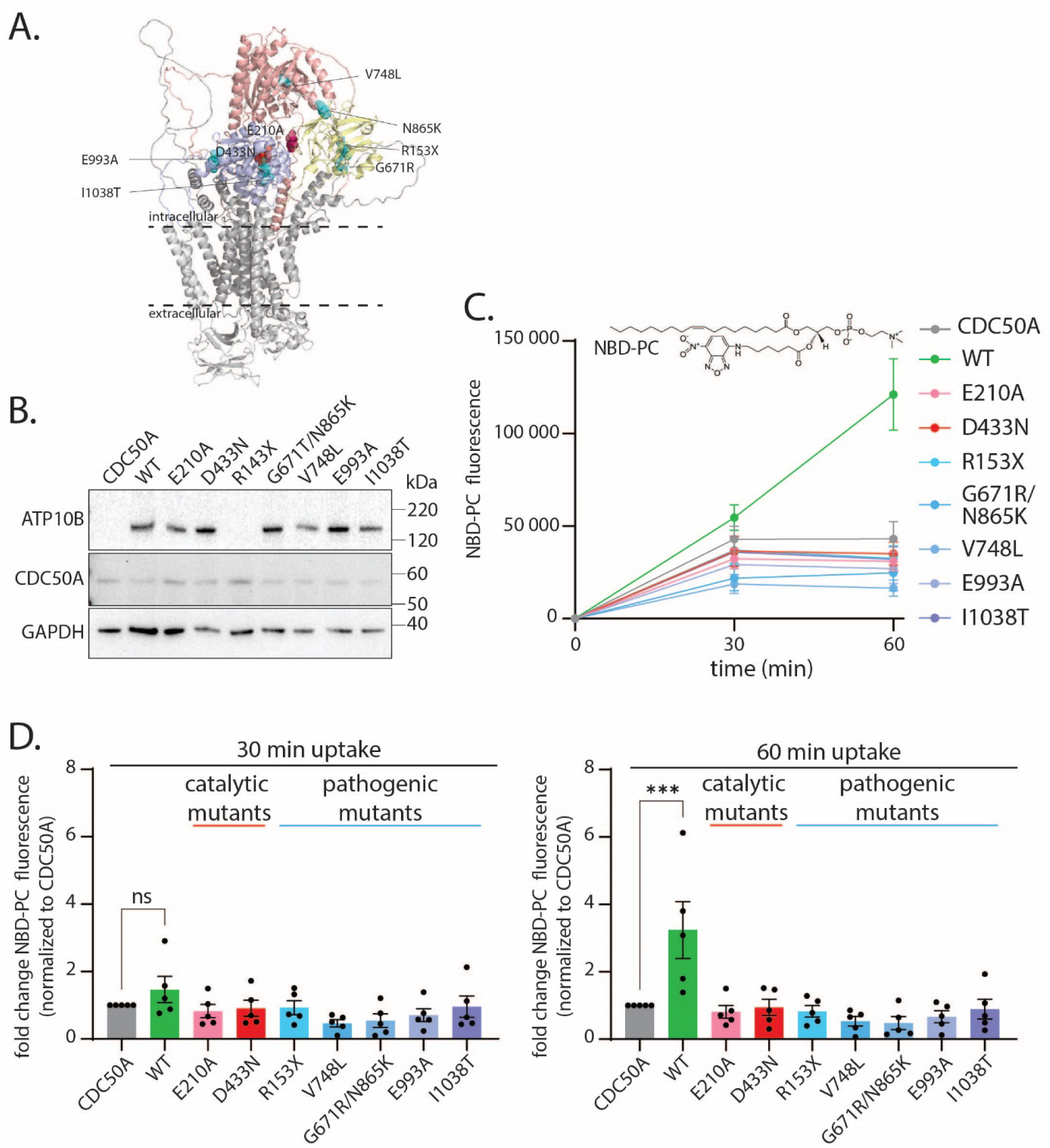
Transiently expressed ATP10B increases cellular PC uptake, while pathogenic variants impair this uptake. **A)** Predicted structure of ATP10B (AlphaFold), P-domain in blue, A-domain in yellow and N-domain in red. Mutations used in this study are shown as spheres (catalytic mutations in red, pathogenic variants in teal). **B)** Representative immunoblot of HeLa cells after transient transfection with CDC50A in combination with ATP10B (WT or mutant). GAPDH is used as a loading control. **C)** Time course of NBD-PC lipid uptake in cells expressing the indicated WT or mutant ATP10B. NBD-PC structure is shown at top of the graph. **D)** NBD-PC uptake normalized to control (expressing only CDC50A) after 30 min (left panel) and 60 min (right panel). Data are represented as mean ± s.e.m., n=5, statistical test was an ordinary one-way ANOVA with Dunnett’s post-hoc pair- wise multiple comparisons test. Significance was set at *P<0.05, **P<0.01, ***P<0.001, and ****P<0.0001.

Next, we assessed the cellular uptake of fluorescently labeled lipids with a lipid uptake assay adapted from (Naito *et al*., 2015). Cells were first incubated with nitrobenzoxadiazole (NBD)-labeled PC, which is transported by ATP10B (Martin *et al*., 2020), and non-internalized lipids were quenched with sodium dithionite prior to the detection of the total internal fluorescence via flow cytometry. After 30 min, the NBD-PC uptake in ATP10B wild type (WT) expressing cells was not significantly different from cells expressing only CDC50A. However, after 60 min, ATP10B WT expressing cells displayed a 3-fold higher NBD-PC uptake compared to CDC50A control cells or the catalytically inactive mutants E210A and D433N (Fig. 1D). The pathogenic protein variants were also unable to increase the cellular NBD-PC uptake (Martin *et al*., 2020). To further confirm the specificity of ATP10B towards NBD-PC, we tested the uptake of other NBD-labeled lipids that based on lipid translocation assays were previously demonstrated not to be substrates of ATP10B (NBD-PE, NBD-PS and NBD-sphingomyelin (NBD-SM)) (Martin *et al*., 2020). The presence of ATP10B WT did not lead to a higher uptake of these NBD-lipids as compared to cells expressing CDC50A or the D433N mutant ATP10B (Supp. 1A-C), demonstrating that the increased cellular uptake is only observed with a biochemically confirmed transported substrate of ATP10B (Martin *et al*., 2020).

### ATP10B-mediated NBD-PC uptake is mediated by stress

To avoid variation in expression levels between independent transient transfections and independently confirm that cellular uptake of NBD-PC is mediated by the endo-/lysosomal lipid flippase ATP10B, we generated stable cell lines with ATP10B and CDC50A overexpression via sequential transduction with CDC50A, followed by ATP10B (WT, catalytic and disease mutants). Viral vectors were titrated to obtain similar expression levels for all ATP10B variants used, which was confirmed via Western blotting (Fig. 2A).

**Figure 2:**
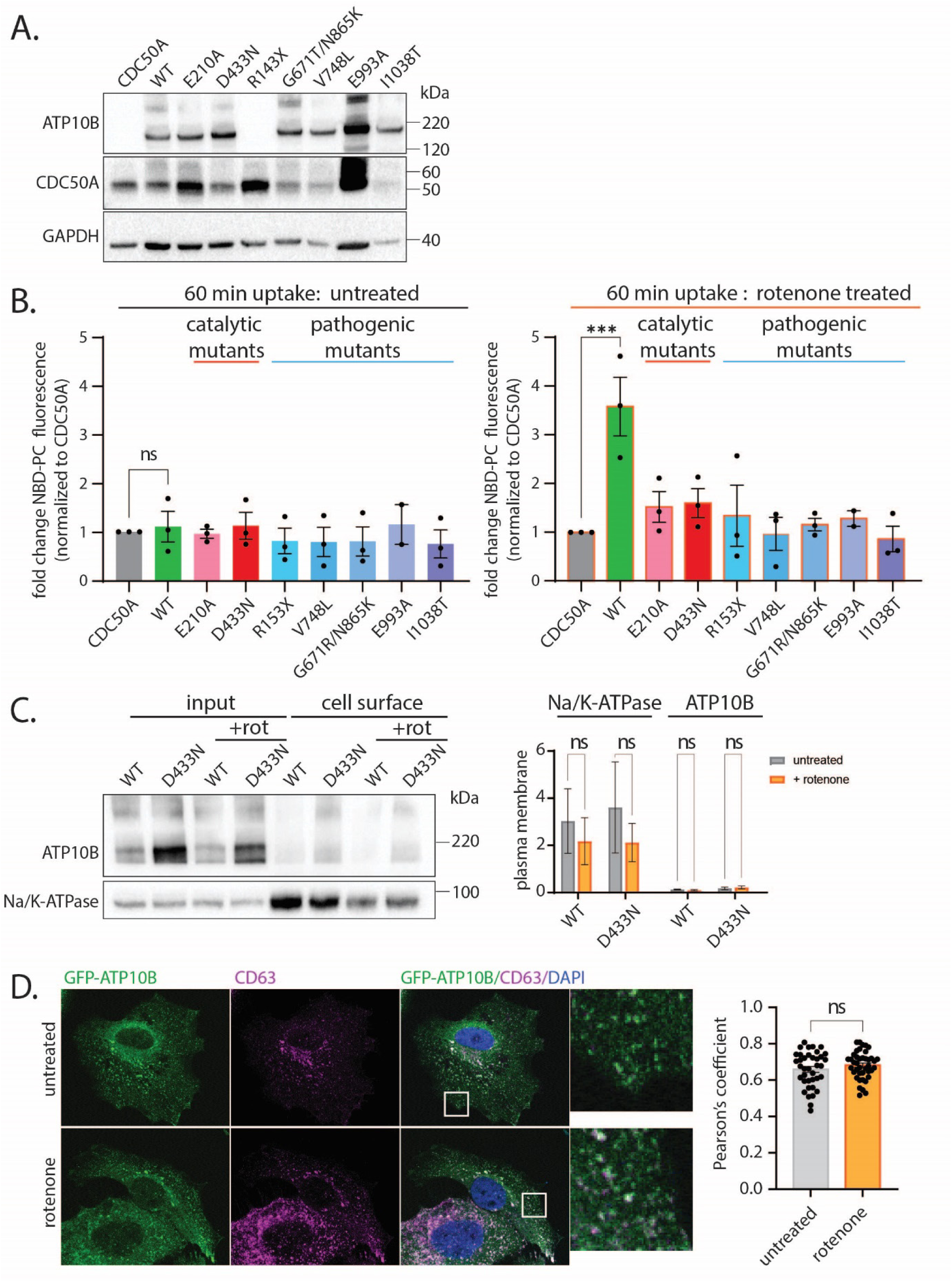
Stably expressed ATP10B increases cellular NBD-PC uptake after 24 h rotenone treatment, not in basal conditions. **A)** Representative immunoblot of HeLa cells stably expressing CDC50A in combination with ATP10B (WT or mutant). GAPDH is shown as a loading control. **B)** NBD-PC uptake normalized to CDC50A control after 60 min of uptake. Uptake was performed on untreated cells (left panel) and cells treated for 24 h with 1 µM rotenone (right panel). **C)** Representative immunoblot and quantification of cell surface biotinylation of cells expressing ATP10B (WT or D433N mutant). Na/K-ATPase is used as positive control. **D)** Colocalization of ATP10B-GFP with endolysosomal marker CD63. Quantification using Pearson’s coefficient. Data are represented as mean ± s.e.m., n=3, statistical test for multiple comparisons (B,C) was an ordinary one-way ANOVA with Dunnett’s post-hoc pair-wise multiple comparisons test, for single comparisons (D) was unpaired t test. Significance was set at *P<0.05, **P<0.01, ***P<0.001, and ****P<0.0001.

Surprisingly, under identical conditions as for the transiently transfected cells, we observed no increase in internalized NBD-PC in the stable cell lines up to 60 min (Fig. 2B, left panel). We therefore hypothesized that ATP10B may reside in an inactive state in the stable cells, in contrast to the transiently transfected cells. Since the transport activity of ATP10B provides cellular protection against the mitochondrial toxin rotenone, a known environmental risk factor for PD, we examined whether rotenone could stimulate ATP10B-mediated NBD-PC uptake in the stable cell lines. Interestingly, after treatment with 1 µM rotenone for 24 h, cells expressing ATP10B WT presented a 3-fold higher NBD-PC uptake as compared to cells expressing the catalytically dead mutants or pathogenic mutants, confirming the ATP10B-mediated uptake of NBD-PC (Fig. 2B, right panel).

Besides the phospholipid PC, the sphingolipid GluCer was previously identified as a substrate of ATP10B (Martin *et al*., 2020). However, in contrast to NBD-PC uptake, ATP10B-mediated NBD-GluCer uptake was not consistently observed between independent experiments (Supp. 2A).

To determine whether the increased NBD-PC uptake under rotenone stress might be caused by the relocalization of ATP10B to the PM, we performed cell surface biotinylation and confocal microscopy to follow ATP10B at the PM and intracellularly. While the known PM protein Na/K-ATPase was clearly detected at the PM by cell surface biotinylation, only small traces of WT or D433N ATP10B were detected (Fig. 2C). While treatment with rotenone moderately, albeit not significantly, decreased Na/K-ATPase levels at the PM, the levels of ATP10B at the PM remained unchanged in rotenone conditions. Additionally, we verified the subcellular localization of ATP10B tagged with GFP, stably expressed in HeLa cells with CDC50A co-expression. The majority of cells presented a clear colocalization of ATP10B-GFP with the late endo-/lysosomal compartment marker CD63, which was unchanged after rotenone treatment (Fig. 2D). A small proportion of cells (<10%), mainly those with high overexpression levels of ATP10B-GFP, did show PM localization of ATP10B-GFP, but these levels did not increase in rotenone conditions (Supp. 2B).

### Endogenous ATP10B regulates cellular NBD-PC uptake

Next, we questioned whether endogenous ATP10B also affects NBD-PC lipid uptake. Since ATP10B is not endogenously expressed in commonly used cell lines, such as HeLa cells, we turned to Im95m cells, a human gastric cancer line with endogenous ATP10B expression. We previously generated two stable shRNA-mediated ATP10B knockdown (KD) lines (Martin *et al*., 2020) with >70% knockdown efficiency at the protein level (Fig. 3A). We then compared NBD-PC uptake in these cells both with and without rotenone stress, and observed a significant difference in uptake between the WT and KD cells, already without rotenone treatment, with the knockdown cells showing a 60% decrease in NBD-PC uptake at 60 min uptake (Fig. 3B). Interestingly, treatment with rotenone further increased the difference to 80% between WT and KD cells for 60 min of uptake (Fig. 3C). This could be recapitulated in human melanoma WM115 cells with KD of endogenous ATP10B, which resulted in a 50% decrease of NBD-PC uptake (Supp. 2).

**Figure 3:**
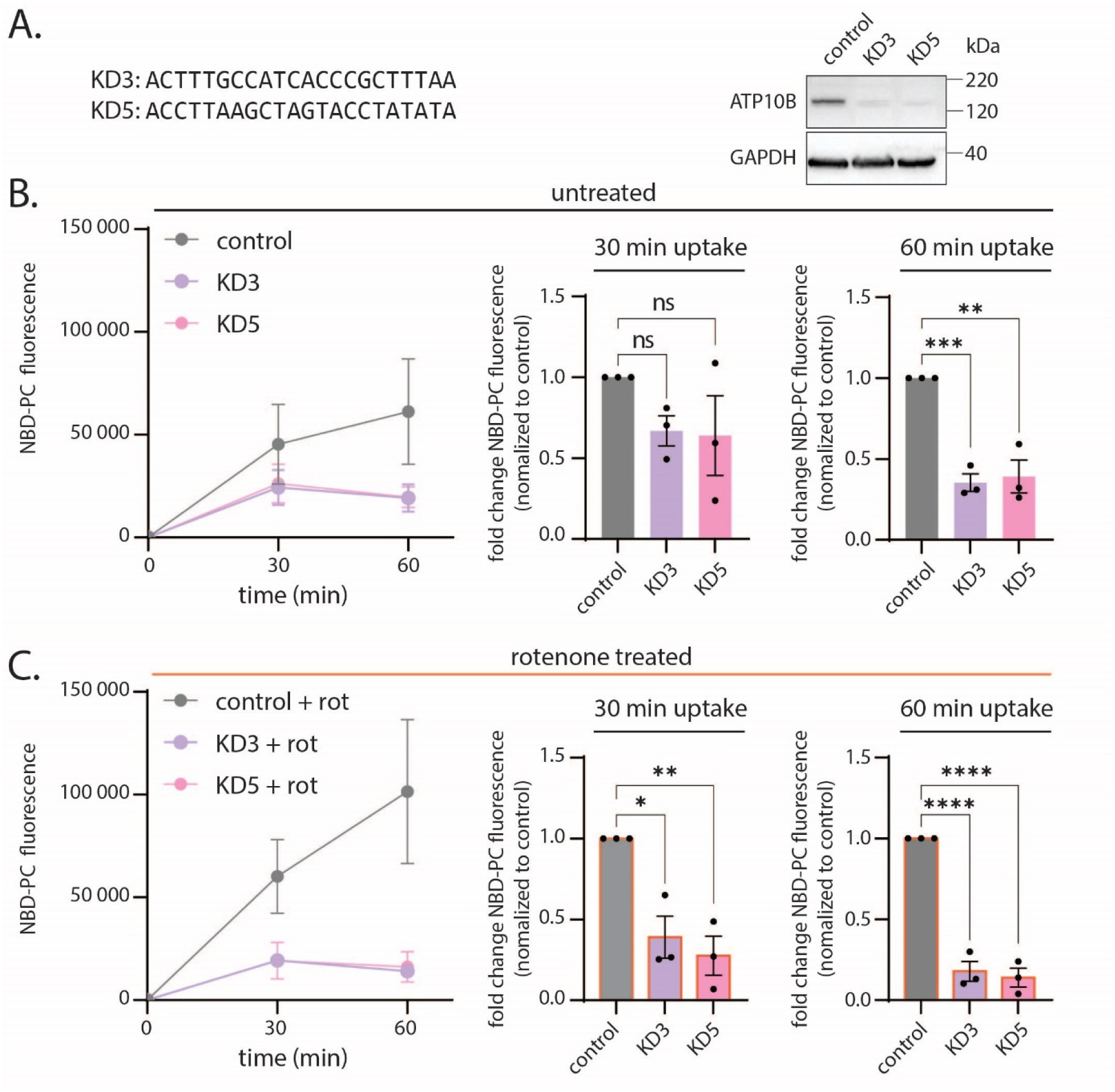
Knockdown of ATP10B in Im95m cells reduces NBD-PC uptake, an effect that is more pronounced after rotenone uptake. **A)** Representative immunoblot of Im95m cells after stable knockdown of endogenous ATP10B with two different miRNA sequences (KD1, KD2). GAPDH is used as a loading control. **B-C)** NBD-PC uptake without (B) and with (C) 24 h rotenone treatment (1 µM). Time course (left panel), uptake normalized to control (expressing only CDC50A) after 30 min (middle panel) and 60 min (right panel). Data are represented as mean ± s.e.m., n=3, statistical test was an ordinary one-way ANOVA with Dunnett’s post-hoc pair-wise multiple comparisons test. Significance was set at *P<0.05, **P<0.01, ***P<0.001, and ****P<0.0001.

Finally, we examined whether ATP10B may affect the endogenous lipid homeostasis and performed a lipidomics analysis on the overexpression and knockdown cells. In both basal and rotenone treated conditions, we found no significant changes in PC levels in HeLa overexpressing ATP10B WT compared to D433N, nor in WM115 KD cells (Supp. 4). Other phospholipids, such as PS and PI, were also unchanged. However, HeLa cells with ATP10B WT expression showed a significant decrease in hexosylceramide (HexCer, *i.e.* the combined lipid pool of GluCer and galactosylceramide (GalCer)), but no changes were observed in the WM115 KD model.

## Discussion

In complementary overexpression and knockdown cell models, we demonstrated that overexpression of ATP10B increases, whereas KD of endogenous ATP10B decreases, uptake of NBD-PC in cells. Since disease variants abrogate the lipid uptake phenotype, this defect possibly contributes to neuronal dysfunction in early onset PD or DLB patients carrying ATP10B variants.

### ATP10B stimulates lipid uptake

We demonstrated that ATP10B-mediated lipid uptake depends on its transport activity, and is selective for a biochemically confirmed transported substrate of ATP10B (Martin *et al*., 2020). We further observed that the ATP10B-mediated lipid uptake depends on cell stress conditions (transient transfection or rotenone treatment). P4-Type ATPase are subjected to auto-inhibition by N- and/or C-terminal extensions, and transport activation typically requires phosphorylation by protein kinases or binding of certain regulatory lipids like phosphatidylinositol lipids (Timcenko *et al*., 2019; López-Marqués *et al*., 2020). Unfortunately, the precise regulatory elements to activate ATP10B are not yet established, but comparison of post-translational modifications in basal and rotenone conditions may be helpful in the future.

Several possible scenarios may explain how an endo-/lysosomal localized flippase contributes to cellular lipid uptake from the medium. A first option is that a fraction of the ATP10B population may be active at the PM working similar as other PM-localized flippases like ATP10A and ATP10D. In 10% of the cells that exhibit the highest expression levels, we indeed observed traces of ATP10B-GFP at the cell surface (Supp. 2B). An imbalance between the levels of ATP10B and the CDC50A, which is required for correct localization, could be responsible for this (Takatsu *et al*., 2011; Naito *et al*., 2015). However, this PM-localized fraction did not alter with rotenone treatment despite an increased NBD-PC uptake in these conditions (Fig. 2D), indicating that the mere presence of ATP10B in the PM does not explain the phenotype. In addition, we found that ATP10B KD also impacted on the NBD-PC uptake (Fig. 3) indicating that it is not an artefact of the overexpression. It may be possible that the minor ATP10B pool at the plasma membrane becomes activated via post-translational modifications, lipid or protein interactions, stimulating NBD-PC uptake (Fig. 4).

**Figure 4:**
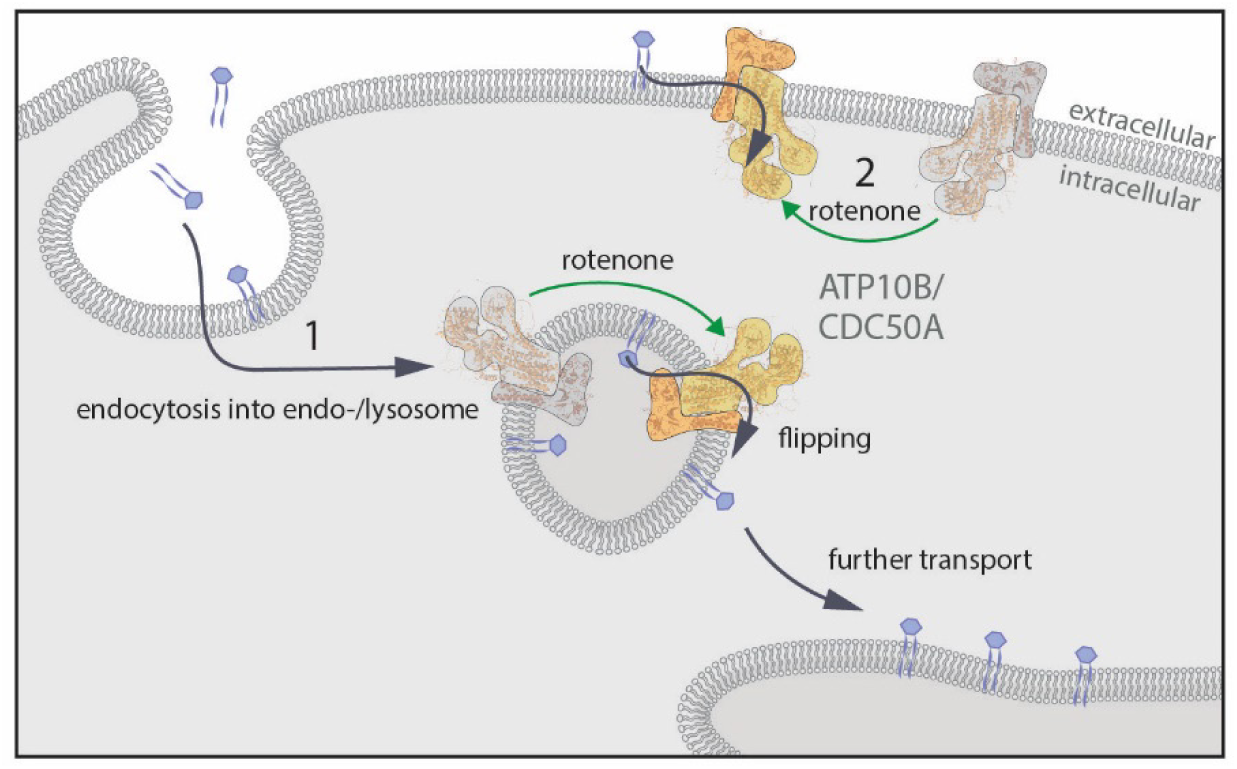
Model of ATP10B function in lipid uptake. Under certain stress conditions, such as rotenone treatment, ATP10B mediates cellular lipid uptake, most likely by flipping its substrate lipids from the exoplasmic to the cytoplasmic membrane leaflet. 1. The majority of ATP10B resides in endo/lysosomal compartments, which may enable cellular lipid uptake via endocytosis prior to transport by ATP10B. 2. A small fraction of the cells presents ATP10B at the plasma membrane, which following activation in rotenone conditions, may contribute to cellular lipid uptake from within the plasma membrane.

A second explanation for our results is that ATP10B from within the late endo-/lysosomes may contribute to NBD-PC uptake via endocytosis (Fig. 4). Such a mechanism would resemble how another late endo-/lysosomal P-type ATPase ATP13A2 stimulates cellular polyamine uptake, which first involves endocytosis prior to late endo-/lysosomal export into the cytosol (van Veen *et al*., 2020). Indeed, matching previous reports, the majority of ATP10B in our overexpression system is present in the endo-/lysosomal system, as shown here through a combination of cell surface biotinylation and imaging (Fig. 2) (Martin *et al*., 2020; Okamoto *et al*., 2020), which fits also with the endosomal targeting motifs previously identified in ATP10B (Takatsu *et al*., 2011). The ATP10B-dependent NBD-PC uptake was seen in the timeframe between 30 min and 60 min of uptake, which takes longer than for a PM-localized lipid flippases that causes near-instantaneous lipid uptake (Naito *et al*., 2015). In the first 30 min, uptake levels were similar between cells expressing WT or catalytically dead variant D433N, suggesting that in this time span other PM-localized lipid flippases are playing a role (Fig. 1C-D). For example, ATP10A is a known PM-localized flippase with PC as substrate that in a HeLa overexpression model stimulates NBD-PC uptake already after 5 min incubation (Naito *et al*., 2015). In this scenario, ATP10B activation would take place in the endosomal compartments. Finally, it remains possible that a change in the endocytosis rate may be responsible for the altered uptake of NBD-PC in ATP10B cell models. However, the process appears selective for NBD-PC uptake, and requires ATP10B transport activity, suggesting that altered endocytosis alone may not be the only reason.

### ATP10B selectively stimulates cellular uptake of PC

While we observed that ATP10B reproducibly stimulates the uptake of NBD-PC (Fig. 1C-D; Fig. 2C), the results with NBD-GluCer were more variable. NBD-GluCer uptake in ATP10B overexpressing cells was only observed in a minority of experiments (Supp. 2A). In previous biochemical experiments, both PC and GluCer were identified as substrates for ATP10B ATPase activity (Martin *et al*., 2020), and also translocation of NBD-PC and NBD-GluCer via ATP10B has been observed indicating that both fluorescent lipids are genuine substrates of ATP10B. The differences between the biochemical and cellular uptake assays may therefore be related to the cellular uptake pathways for both lipid species. The uptake of sphingolipids may involve caveolar endocytosis (Shvets *et al*., 2015; Zhou *et al*., 2021), whereas NBD-PC may largely depend on clathrin-dependent endocytosis (Singh *et al*., 2003). We hypothesize that PC, as a principal component of most membranes (Van Meer, Voelker and Feigenson, 2008), reaches ATP10B through parallel endocytosis routes, whereas GluCer may rely on more dedicated uptake pathways for sphingolipids that not necessarily end up in the late endo-/lysosomal compartment. Indeed, it has been proposed that sphingolipids may be mainly transported from endosomes to the Golgi, only reaching lysosomes in sphingolipid storage disorders (Puri *et al*., 2001). GluCer also forms microdomains within membranes, in a pH-dependent manner, which may alter the membrane organization during the maturation process of endo-/lysosomal vesicles (Varela *et al*., 2016). The distribution of ATP10B within these lipid domains will determine its access to lipid substrates for transport.

Surprisingly, the effect of ATP10B on NBD-PC *versus* NBD-GluCer uptake does not correlate well with changes in the total lipid levels (Supp. 4). While there is a clear effect of ATP10B on NBD-PC uptake (Fig. 1C-D; Fig. 2C), but not on NBD-GluCer uptake (Supp. 2A), total PC levels remain unchanged (Supp. 4A-B), whereas HexCer levels alter only in an overexpression model (Supp. 4A). These differences might be explained in the light of the different experimental conditions during the assays. The sudden and transient addition of exogenous NBD-labeled lipids during the uptake experiments shows a rapid uptake of lipids following lipid supplementation. For the lipidomics analysis, cells have not been challenged by lipid supplementation, and therefore, our results reflect basal lipid homeostasis, which is a combination of lipid import, export and metabolism. Since many factors contribute to regulate the cellular lipid levels, it is possible that ATP10B modulation goes hand in hand with differences in lipid metabolism and transport pathways that buffer changes in lipid uptake. PC, as a principal component of many cellular membranes, might follow many parallel transport routes including ATP10B, which may explain a relatively smaller impact of ATP10B-mediated PC transport on PC levels. On the other hand, fewer transporters are known for GluCer, suggesting that the relative impact of ATP10B on total GluCer levels may therefore be larger. Additionally, since the relative concentration of GluCer compared to PC in membranes is lower, ATP10B activity may have a stronger impact on GluCer levels. While our results indicate that ATP10B stimulates the endogenous HexCer content, the relative contribution of uptake versus metabolism may depend on the cell type and stress context (Else, 2020). To further evaluate the role of ATP10B in the context of PD, we plan further experiments in iPSC and animal models to confirm the physiological impact of ATP10B on PC and GluCer homeostasis.

## Conclusion

We demonstrated that the late endo-/lysosomal ATP10B stimulates cellular PC uptake, which is dependent of the cellular conditions. Conversely, ATP10B mainly impacts the endogenous cellular HexCer levels, which appears cell type specific. Our results indicate that studies in PD-relevant models will be required to firmly establish the physiological role of ATP10B in lipid homeostasis.

## Acknowledgements

This research was funded by the Aligning Science Across Parkinson’s [ASAP-000458] through the Michael J. Fox Foundation for Parkinson’s Research (MJFF) allocated to PV and VB. The research was also supported by a C1 KU Leuven grant (C15/15/073), the Fonds voor Wetenschappelijk Onderzoek (FWO) Flanders (S006617N SBO Neuro-Traffic, PV and VB), Michael J. Fox Foundation MJFF-008610 (with generous support of the Demoucelle Parkinson Charity and the Stop Parkinson Walk) (PV) and the Queen Elisabeth Medical Foundation for Neurosciences Valine de Spoelberch award (PV).

## Abbreviations

BSA: bovine serum albumin
FBS: fetal bovine serum
GalCer: galactosylceramide
GCase: glucocerebrosidase
GluCer: glucosylceramide
HBSS: Hanks balanced sodium solution
HexCer: hexosylceramide
KD: knockdown
NEEA: non essential amino acids
NBD: nitrobenzoxadiazole
PBS: phohsphate buffered saline
PC: phosphatidylcholine
PD: Parkinson’s disease
PE: phosphatidylethanolamine
PM: plasma membrane
PS: phosphatidylserine
SM: sphingomyelin
WT: wild type

---

**Supplementary Figure 1:**
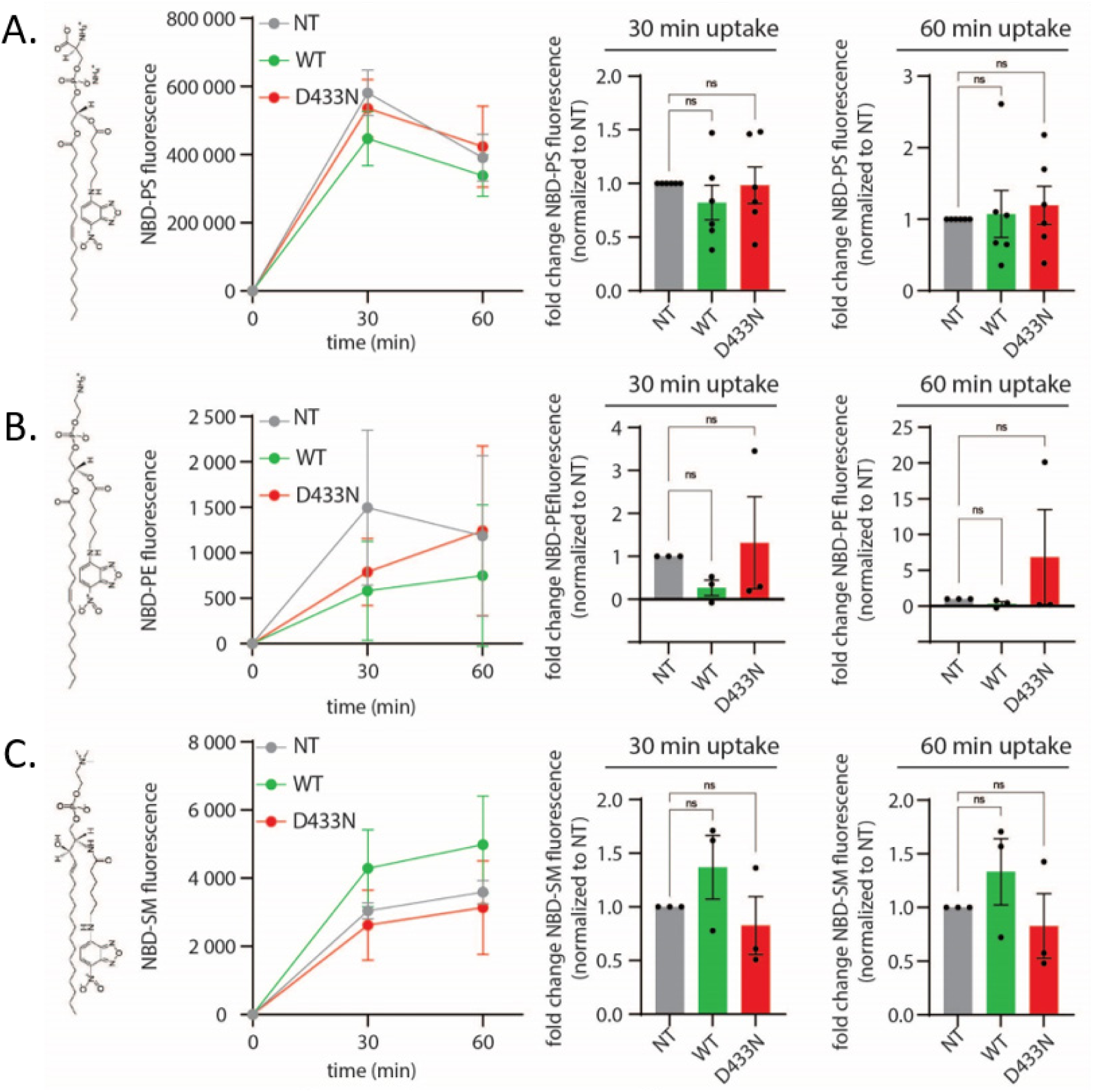
ATP10B-mediated lipid uptake is specific for NBD-PC. **A-C)** NBD-lipid uptake after no transient transfection or combined transient transfection with ATP10B WT or ATP10B D433N. Lipids used were (A) NBD-PS, (B) NBD-PE, and (C) NBD-SM. Time course (left panel), uptake normalized to control (expressing only CDC50A) after 30 min (middle panel) and 60 min (right panel). Data are represented as mean ± s.e.m., n=3, statistical test was an ordinary one-way ANOVA with Dunnett’s post-hoc pair-wise multiple comparisons test. Significance was set at *P<0.05, **P<0.01, ***P<0.001, and ****P<0.0001.

**Supplementary Figure 2:**
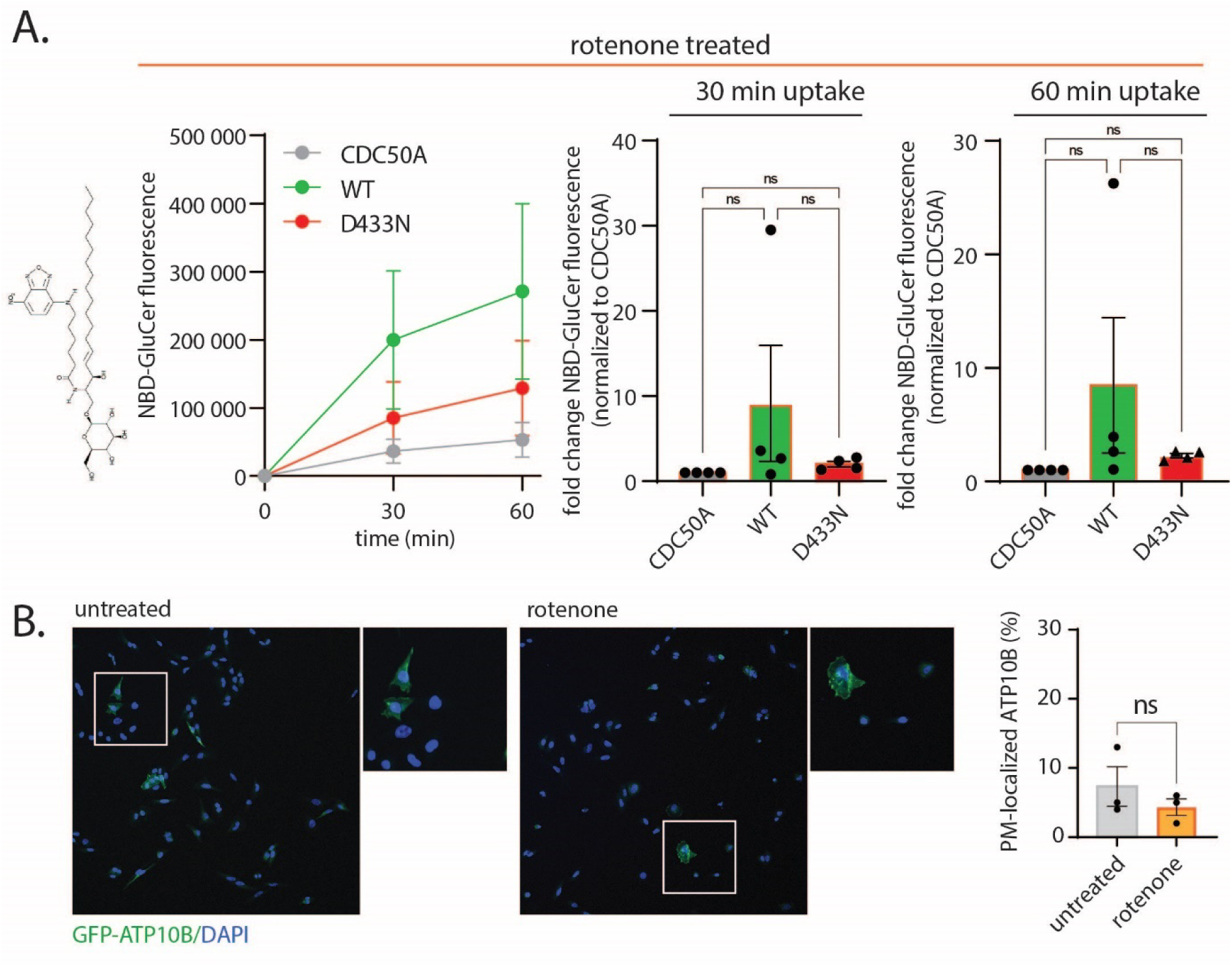
Rotenone treatment does not induce ATP10B-mediated uptake of NBD- GluCer or redistribute ATP10B-GFP to the plasma membrane. **A)** NBD-GluCer uptake in HeLa cells with stable expression of CDC50A alone or in combination with ATP10B WT or ATP10B D433N, after 24 h rotenone treatment (1 µM). Time course (left panel), uptake normalized to control (expressing only CDC50A) after 30 min (middle panel) and 60 min (right panel). **B)** Confocal imaging of GFP- ATP10B stably expressed in HeLa cells. Percentage of cell with GFP-ATP10B localized at the plasma membrane was quantified. Data are represented as mean ± s.e.m., n=4 (A) or n=3 (B), statistical test for multiple comparisons (A) was an ordinary one-way ANOVA with Dunnett’s post-hoc pair-wise multiple comparisons test, for single comparisons (B) was unpaired t test. Significance was set at *P<0.05, **P<0.01, ***P<0.001, and ****P<0.0001.

**Supplementary figure 3:**
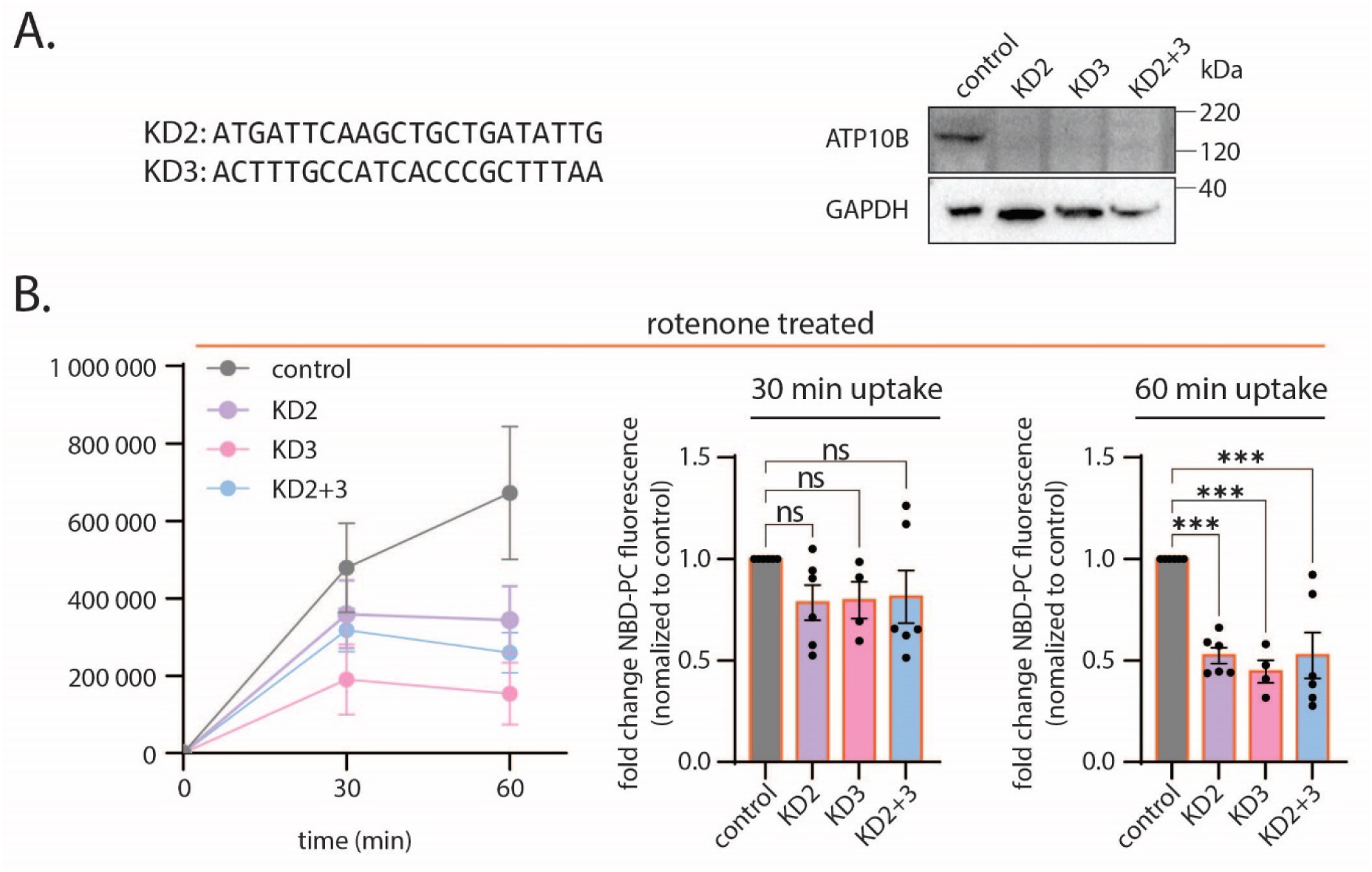
Knockdown of ATP10B in WM115 cells decreases NBD-PC uptake after rotenone treatment. **A)** Representative immunoblot of WM115 cells after stable knockdown of endogenous ATP10B with two different miRNA sequences separately (KD2, KD3) or combined (KD2+3). GAPDH is used as a loading control. **B)** NBD-PC uptake with 24 h rotenone treatment (1 µM). Time course (left panel), uptake normalized to control (expressing only CDC50A) after 30 min (middle panel) and 60 min (right panel). Data are represented as mean ± s.e.m., n=6 (control, KD2, KD2+3) or n=4 (KD3) (B), statistical test for multiple comparisons was an ordinary one-way ANOVA with Dunnett’s post-hoc pair-wise multiple comparisons test. Significance was set at *P<0.05, **P<0.01, ***P<0.001, and ****P<0.0001.

**Supplementary figure 4:**
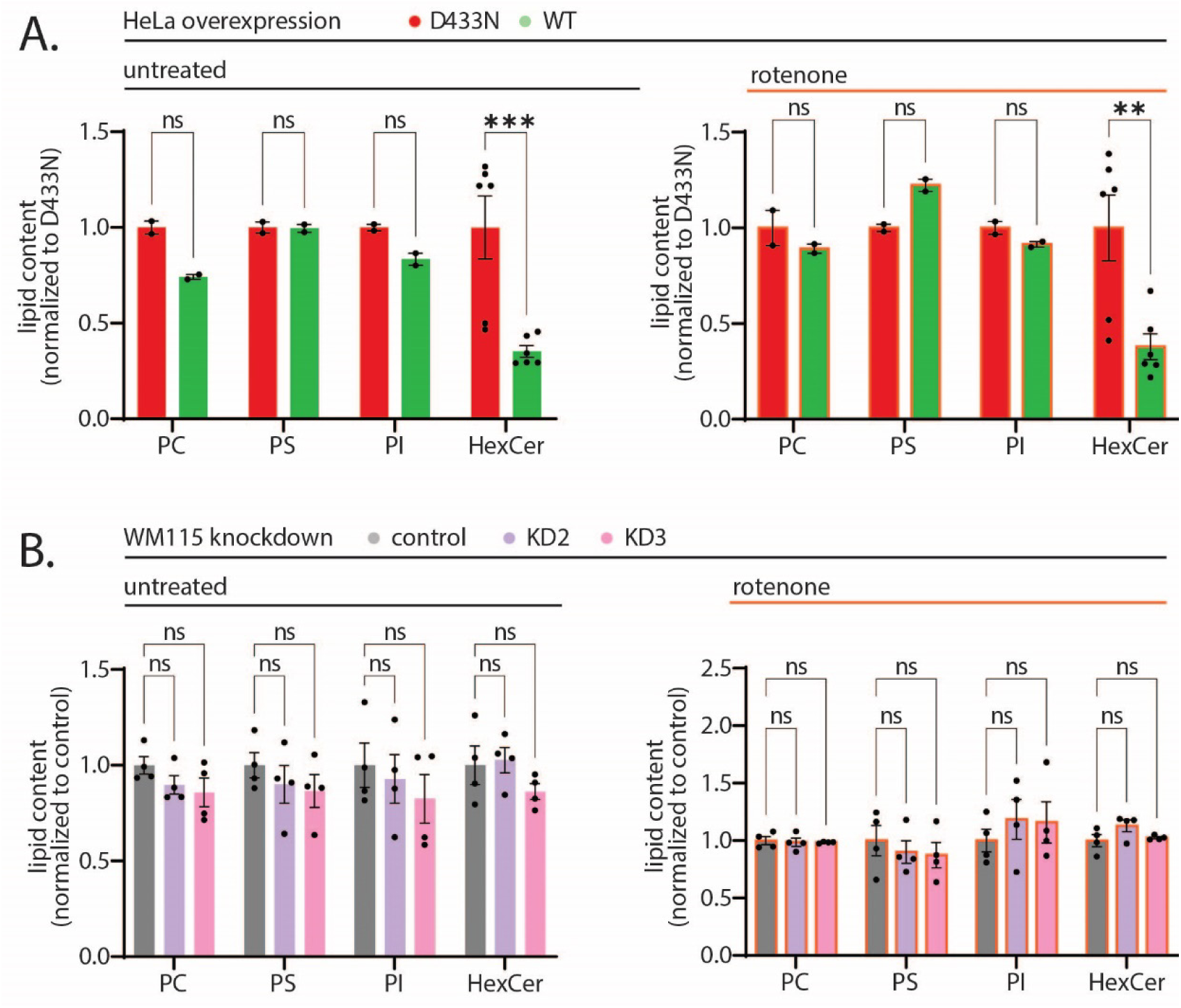
Total cellular lipid levels can be affected by ATP10B expression levels. **A)** Lipidomics of HeLa with stable expression of CDC50A in combination with either ATP10B WT or D433N. PC, PS and PI are measured as nmol lipid/mg DNA, HexCer is measured as analyte/internal standard. **B)** Lipidomics of WM115 cells, comparing control to ATP10B KD. All lipids are measured as nmol/mg DNA. Data are represented as mean ± s.e.m., n=2 (A (PC, PS, PI)), n=6 (A (HexCer)), n=4 (B), statistical test was an ordinary one-way ANOVA with Dunnett’s post-hoc pair-wise multiple comparisons test. Significance was set at *P<0.05, **P<0.01, ***P<0.001, and ****P<0.0001.

